# Multimodal magnetic resonance imaging predicts regional amyloid- β burden in the brain

**DOI:** 10.1101/2020.01.17.910984

**Authors:** Anusha Rangarajan, Minjie Wu, Naomi Joseph, Helmet T. Karim, Charles Laymon, Dana Tudorascu, Annie Cohen, William Klunk, Howard J. Aizenstein

**Affiliations:** Departments of Bioengineering, University of Pittsburgh, Pittsburgh, PA, USA; Departments of Psychiatry, University of Pittsburgh, Pittsburgh, PA, USA; Departments of Chemical Engineering, University of Pittsburgh, Pittsburgh, PA, USA; Departments of Radiology, University of Pittsburgh, Pittsburgh, PA, USA; Departments of Biostatistics, University of Pittsburgh, Pittsburgh, PA, USA; Departments of Medicine, University of Pittsburgh, Pittsburgh, PA, USA

**Keywords:** amyloid- β, prediction, multimodal MRI, regression, Alzheimer’s Disease

## Abstract

Alzheimer’s disease (AD) is the most common cause of dementia and identifying early markers of this disease is important for prevention and treatment strategies. Amyloid - *β* protein deposition is one of the earliest detectable pathological changes in AD. But in-vivo detection of amyloid - *β* using positron emission tomography (PET) is hampered by high cost and limited geographical accessibility. These factors can become limiting when PET is used to screen large numbers of subjects into prevention trials when only a minority are expected to be amyloid- β - positive. Structural MRI is advantageous; as it is relatively inexpensive and more accessible. Thus it could be widely used in large studies, even when frequent or repetitive imaging is necessary. We used a machine learning, pattern recognition, approach using intensity-based features from individual and combination of MR modalities (T1 weighted, T2 weighted, T2 fluid attenuated inversion recovery [FLAIR], susceptibility weighted imaging) to predict voxel-level amyloid- β in the brain. The MR- amyloid *β* relation was learned within each subject and generalized across subjects using subject–specific features (demographic, clinical, and summary MR features). When compared to other modalities, combination of T1-weighted, T2-weighted FLAIR, and SWI performed best in predicting the amyloid- β status as positive or negative. T2- weighted performed the best in predicting change in amyloid- β over two timepoints. Overall, our results show feasibility of amyloid- β prediction by MRI.

## 1. Introduction

Alzheimer’s disease (AD) is a neurodegenerative disease affecting more than 5 million people in the US. It has been identified as the most common cause of dementia in people 60 years and older (Alzheimer’s, 2019). The disease is believed to begin with pathological changes, including accumulation of amyloid- β protein; followed by tau protein tangles, hypometabolism, inflammation and brain atrophy several years before the cognitive and clinical symptoms are apparent (Jack et al., 2010; Sperling et al., 2011). As one of the earliest detectable pathological changes in AD, amyloid- β deposition is a primary target for prevention and treatment strategies (Weninger et al., 2016). In-vivo measurement of amyloid- β plaques is performed using positron emission tomography (PET)(Villemagne, 2016). There is evidence that biormarkers identifiable by PET (Amyloid- β and hypo-metabolism) appear during the preclinical stages of AD (Cohen and Klunk, 2014; Jack et al., 2012; Marcus et al., 2014; Sperling et al., 2011) and can predict decline years before the onset of symptoms (Cohen and Klunk, 2014). In-vivo amyloid- β imaging is essential in definitive diagnosis of AD and important for the early detection of AD (Adlard et al., 2014). However limited access and high cost restrict the use of amyloid- β PET ‘ scans.

Structural MRI is widely accessible and is relatively inexpensive. If MRI predictors of amyloid- β deposition could be defined, MRI could have advantages over PET in several areas: as a screening tool in large prevention studies where large numbers of potential subjects need to be screened, where frequent or repetitive imaging is necessary, and perhaps for monitoring disease progression and evaluation of treatment.

Greater levels of amyloid- β plaque in the brain are associated with worsened structural brain integrity. Amyloid-β deposition is associated with reduced cortical thickness in parietal, posterior cingulate and precuneus regions (Becker, 2011) (Dore, 2013) and hippocampus (Dore, 2013) more specifically entorhinal cortex (Doherty, 2015). Regional changes in brain structure quantified using changes in MR intensities can help predict regional amyloid- β deposition. The subtle changes in the local pattern of image intensity (textures) across voxels can be detected using image texture analysis (Maani et al., 2015).

MRI texture analysis studies have shown promise in characterizing brain tumors (Zacharaki et al., 2009) (Bahadure et al., 2017), multiple sclerosis (Abbasian Ardakani et al., 2015) (Harrison et al., 2010) and epileptic seizure prediction (Suoranta et al., 2013) (de Oliveira et al., 2013). Quantitatively image textures can be captured using various spatial or frequency based filters. There are multiple MR modalities that highlight different tissue properties in the brain. High-resolution T1-weighted images have shown great tissue contrasts from which summary measures like volume and thickness measurements and texture-based measures are obtained. T2-weighted imaging can help detect fluid accumulation in the brain (Vemuri et al., 2017). T2-weighted FLAIR helps visualize white matter lesions (WMLs) as hyperintensities (Moller et al.). These white matter hyperintensities (WMHs) are highly correlated with AD (Brickman et al., 2009; Kandel et al., 2016) and amyloid-β deposition (Noh et al., 2014; Park et al., 2014). Tissue magnetic susceptibility differences highlighted using susceptibility weighted imaging (Moller et al.) help visualize microbleeds. Clinically SWI has been used for visualizing vasculature in the brain which show T2* differences in blood and surrounding tissues ((Di Ieva et al., 2015) (Hsu et al., 2017),(Liu et al., 2017),(Halefoglu and Yousem, 2018)). These microbleeds are also associated with AD (Sepehry et al., 2016) and amyloid- β (Graff-Radford et al., 2018)(Dierksen, 2010).

In this study, we use intensity based voxel-level imaging features extracted from both individual modalities and a combination of modalities to predict voxel-level amyloid- β. We use regression models to perform amyloid- β status prediction across subjects (least absolute shrinkage and selection operator (LASSO) and partial least squares (PLS)) and amyloid- β change prediction within subjects (LASSO). Imaging data from fourteen subjects are used in the within subject prediction and thirty-five subjects are used for the prediction across subjects. The machine learning algorithm takes advantage of the voxel-wise data. Thus, the effective sample size for machine learning is > 10,000 examples. The MRI based approach proposed here could offer promise for characterizing voxel-level amyloid- β burden on cross-sectional data and also potentially tracking amyloid- β burden longitudinally.

## 2. Materials and Methods

### 2.1. Parent Study and Subjects

This study was part of an ongoing longitudinal study (started in 2007) (RF1 AG025516) at the University of Pittsburgh. In 2011, the MR scanning for this study switched to a Siemens 3T TRIO MR scanner. For the current study we selected all subjects who underwent at least one scan on the 3T TRIO scanner between 2011 and 2017). If they had more than one session on the 3T scanner then we chose the 1^st^ scan that had all MR sequences (T1-weighted, T2-weighted, SWI, T2 FLAIR). Subjects (for amyloid- β change prediction, N=13 with two time-points; for amyloid- β status prediction across subjects, N=35) were scanned using both MRI and PET on separate visits. Mean age for the subjects were 76 (5.7) years, gender were 62.8% female, race distribution was 6 African American, 1 Asian and 28 White. All subjects signed written informed consent approved by the University of Pittsburgh institutional review board

### 2.2. MRI Acquisition

All MRI scanning was conducted using a 3T Siemens Trio (Munich, Germany) located at the MR Research Center at the University of Pittsburgh with a 12-channel head coil. An axial, whole brain (3D) MPRAGE was collected with echo time (TE)=2.98ms, repetition time (TR)=2300ms, flip angle (FA)=9°, inversion time (TI)=900ms, field of view (FOV)=256×240, 1.2×1×1 mm, and 160 slices. An axial, whole brain (2D) T2-weighted image was collected with TE=101ms, TR=5300ms, FA=150°, TI=2500ms, FOV=256×256, 1×1×3 mm resolution, and 48 slices. An axial, whole brain (2D) FLAIR was collected with TE=90ms, TR=9160ms, FA=150°, TI=2500ms, FOV=256×212, 1×1×3 mm resolution, and 48 slices. An axial, whole brain (2D) SWI was collected with TE=20ms, TR=28ms, FA=15°, TI=300ms, FOV=230×179, 0.53×0.53×1.5 mm resolution, and 96 slices.

### 2.3. PET scanning: Pittsburgh Compound-B (PiB)

[11C]PiB was produced as previously described (Price 2005). PET imaging was conducted using a Siemens/CTI ECAT HR + (3D mode, 15.2 cm field-of-view, 63 planes, reconstructed image resolution ~ 6 mm FWHM). The participant’s head was immobilized to minimize head motion.

PiB was injected intravenously (12–15 mCi, over 20 s, specific activity ~ 1–2 Ci/μmol) and PET scanning was performed over an interval that included 50-70 min post injection. Images were quantified using standard uptake value ratio (ratio of tissue radioactivity to the reference region(cerebellum)).

### 2.4. MRI Preprocessing

All data were preprocessed using statistical parametric mapping software (SPM12). All structural scans (T2-weighted, FLAIR SWI) were linearly registered (normalized mutual information similarity metric and 4^th^ degree B-spline interpolation) to the MPRAGE. These images were then segmented (using the multi-spectral segmentation that utilizes each channel to improve segmentation) into gray matter (GM), white matter (WM), cerebrospinal fluid (CSF), skull, soft-tissue, and air (outputs a probability map for each class). The GM, WM, and CSF were threshold at a probability of 0.1, and added to create an initial intracranial volume mask, which was then refined using an image filling algorithm as well as an image-closing (disk structuring element of 1 voxel) algorithm in MATLAB. The MPRAGE was then linearly registered (normalized mutual information similarity metric with 4^th^ degree B-spline interpolation) to the PET image and that transformation was applied to all other structural scans as well as the GM, WM, and CSF segmentations.

For the longitudinal data an additional within subject registration was performed. The MRI data from two timepoints were registered to the PET data of the baseline. In addition to that the PET data were also registered to the baseline timepoint PET data.

### 2.5. PET Processing

PET data were corrected for photon attenuation, scatter, scanner deadtime, accidental coincidences, and radioactive decay. The final reconstructed PET image resolution was ~6 mm (transverse and axial) based on in-house point source measurements. If subject motion was present then the data were also corrected for inter-frame motion by applying a more extensive registration procedure prior to the PET to MR alignment. The ROI sampling is performed using ROITOOL (Interactive Data Language, Boulder,CO) where the ROIs were all in the PET data space.

There were hand-drawn set of six regions as previously defined (Cohen et al., 2009), which include frontal cortex (FRC; ventral and dorsal), anterior cingulate gyrus (ACG: subgenual and pregenual), anteroventral striatum (AVS), precuneus/posterior cingulate cortex (PRC; ventral, middle and dorsal thirds), parietal cortex (PAR), lateral temporal cortex (LTC), and cerebellum (Bozzali et al.). A global PiB retention index reflecting cerebral amyloid- β is computed from a weighted average of the SUVR values from the six most relevant VOIs (ACG, FRC, LTC, PAR, PRC, and AVS). Subjects were classified as amyloid- β positive or negative based on a global threshold determined from an separate sparse k-means cluster analysis (Cohen et al., 2013).

### 2.6. Overview of Prediction Algorithm

Our overall approach is designed to take advantage of the high dimensionality of the within-subject imaging data to predict the MR-amyloid- β relationship. The MR-amyloid- β relationship developed within subject is used for prediction on longitudinal change in amyloid- β. We then generalize the MR-amyloid- β relationship across subjects, using the subject-level features, to dichotomize amyloid- β status.

#### 2.6.1. Determining voxel –level association between MR and Aβ

*Feature Extraction*

Features were obtained from the native resolution of MR modalities before registration to PET data (i.e., T1, T2, T2 FLAIR and SWI). The specific imaging features used are intensity, 3D gradient, Gabor filter, local binary pattern (intensity-based neighborhood encoding method with detailed description in supplement). Texture can be described as fine or coarse, regular or irregular, and homogeneous or heterogeneous. Gabor filters help characterize homogeneity of texture in the spatial frequency domain. Local binary patterns are a unique way of encoding neighborhood intensity information. 3D gradients highlight regions with rapid change in intensities across voxels. MR signal intensities and Gabor filtered images were obtained from all modalities. 3D gradient features and local binary patterns (LBP) and are usually summarized as histograms. For highlighting voxel-level changes, these features will be most useful in T1- weighted imaging due to its high-resolution with clear contrast in intensities between gray matter and white matter. Hence the voxel-level LBP and 3D gradients are obtained only from T1- weighted imaging. Feature images were then registered to PET (PiB). A combined gray matter and white matter mask (GM+WM) was applied to each of these feature images for obtaining the feature matrix *X_vxF_*; *where F* — *imaging features* (*F* — 17 (*T*1), 13(*T*2, *T*2 — *FLAIR, SWI*)) *and v* — *number of voxels*. The distributions of the features were skewed hence a log_10_ transform was performed.

*LASSO Model*

LASSO is a regularized regression that penalizes the number of non-zero parameter estimates and allows for selection of features that are most predictive of the outcome(Tibshirani, 1996). Features were extracted from voxels within combined gray matter (GM) and white matter (WM) mask (for training LASSO) and 6 volumes of interest (for prediction). For training, voxels from GM+WM mask (~20,000 for each subject) were used to allow space for greater variance in MR features and amyloid- β to improve the learning of the voxel-level MR-amyloid- β relationship. The prediction was performed on voxels within 6 VOIs. The general equation for LASSO regression has a least square minimization term and a penalization term, where *λ* is a regularization parameter, *v* is the total number of voxels, *β*_0_,intercept and *β* vector of slopes.

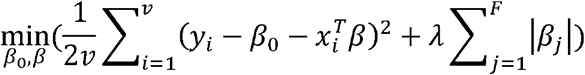

The MR imaging feature matrix, *X_vxF_* (where F are imaging features obtained from MR, *v* are voxels), is used to predict the amyloid- β *y_vx1_*; *where v* — *number of voxels*. The regularization parameter *λ* was estimated via cross-validation using 5% of the voxels. All the voxels within each subject were then used to predict the LASSO parameters *β*_0_,*β*_1_... *β_F_*. These LASSO model parameters can be applied to MR imaging of the same subject at a future time point for predicting amyloid- β change (2.6.1.1). The LASSO models can be generalized across subjects for predicting amyloid- β status (2.6.1.2).

*Evaluation Metrics*

The prediction performance was evaluated using ***accuracy*** (no of. correct label classifications / total number of cases), ***F-score*** defined using ***Precision*** (positive prediction rate) 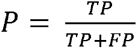 and ***Recall*** 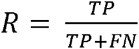 (*where TP* — *true positives, FP* – *false positives, FN* – *false negatives*), area under the receiver operating characteristic (ROC) curve (***AUC***), and ***spearman rank correlation coefficient***. Final individual/combination of modalities are ranked using each of these evaluation metrics and average ranking is used to obtain the most predictive modalities (least rank value).

##### 2.6.1.2. Amyloid- β status prediction across subjects

Subject-level features include demographics (age, weight, sex, race), white matter hyper intensitity burden (WMH), hippocampal volume, and normalized gray and white matter volumes. Partial least squares (PLS) regression was used to learn the relationship between subject-level features and LASSO parameters. Since there are a greater number of amyloid- β negative subjects, we followed the cross-validation scheme established in previous literature (Mathotaarachchi et al., 2017) for rebalancing the data set. We split the data (N=35) into subsets with equal representation of positives and negatives. For each subset, all PiB+ (9) subjects were combined with random combinations of PiB- (9) subjects.

For each subset, a nested leave-one-out-cross-validation (LOOCV)(Browne, 2000) is used for predicting amyloid- β status. An outer LOOCV is performed for testing the amyloid- β status prediction. The inner LOOCV is performed for mapping threshold from original amyloid- β (PiB) range to predicted range (PiB). For each of the test subjects in the outer loop, subject-level features and PLS model are used to predict LASSO parameters. LASSO parameters are applied on voxel-level MR features (within 6 GBL ROIs) to obtain voxel-level amyloid- β (PiB) prediction. The predicted SUVR is range-normalized (range from 0 to 1) and the mean amyloid- β from 6 ROI’s is classified using the mapped threshold as PiB+ or PiB-. An average F-score and AUC across all subsets is used for evaluating the amyloid- β status classification. Additionally, to test whether the subject-level classification accuracy was due to accurate voxel-level PiB prediction, we also tested prediction algorithm using decision trees with only subject level features (i.e., not voxel-wise predicted PiB).

##### 2.6.1.1. Amyloid- β change prediction within subject

Among the total subjects (N=35) used in amyloid- β status prediction, N=13 had two timepoints with all 4 MRI modalities (T1-weighted, T2-weighted, T2-weighted FLAIR, SWI) and PET (PiB). We performed the within subject amyloid- β change prediction using only LASSO regression.

For each subject, LASSO parameters (*β*_0_, *β*_1_... *β_F_*) (*F* – *number of features*) were obtained from MR features and amyloid- β data from time-point 1 and applied on MR imaging features on time-point 2 to predict amyloid- β. Predicted voxel-level amyloid- β deposition can be obtained as a weighted sum of the MR imaging features.

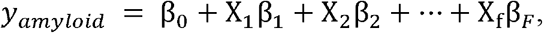

The change in mean SUVR (time-point 2 – time-point 1) was calculated for original and predicted amyloid- β image at the time point. The amyloid- β change detected for all the subjects (N=13) are classified using a threshold obtained at the 50 percentile for both original and predicted change.

## Results

### Amyloid- β status prediction across subjects

The features from multimodal magnetic resonance imaging were able to predict regional amyloid- β. The regional amyloid- β obtained voxel level is then used to calculate the global PiB and that metric is used for classification. The combination of T1-weighted, T2-weighted FLAIR, and SWI modalities performed the best for amyloid- β status classification [Average over 20 subsets: F-score (Mean (SD)): 68.2515 (7.5785), Accuracy (Mean (SD)): 66.3889 (8.156), AUC 0.6639 (0.0816); Spearman correlation coefficient (Mean (SD)): 0.4062 (0.1304)] (Table 1.). Figure 2 shows the mean original and the predicted amyloid- β for one subset (18 subjects) with 3 false positives (FP), 1 false negatives (FN), 8 true positives (TP), and 5 true negatives (TN), thus showing a high sensitivity and medium specificity performance. The false positive predictions could also be an identification of risk subjects. When we examined the 3 false positive subjects we noticed that 2 of them demonstrated regional amyloid- β positivity (one or more of the 6 regions showed high amyloid- β deposition). The one false negative had an original threshold as 1.53 slightly above the threshold (of 1.51) which could be possible that the subject is misclassified. The threshold is predicted for each left-out subject using the nested LOOCV. We found that summary MR measures, demographics, and the combination all performed worse (with decision tree learning) than the voxel-level amyloid- β prediction results described above. The F-scores were less than the predicted amyloid- β from our method [F-score (Mean (SD)) (20 subsets): demographics only = 0.55(0.16), MR only = 0.57(0.16), Demographics and MR = 0.54(0.18)].

**Figure 1:**
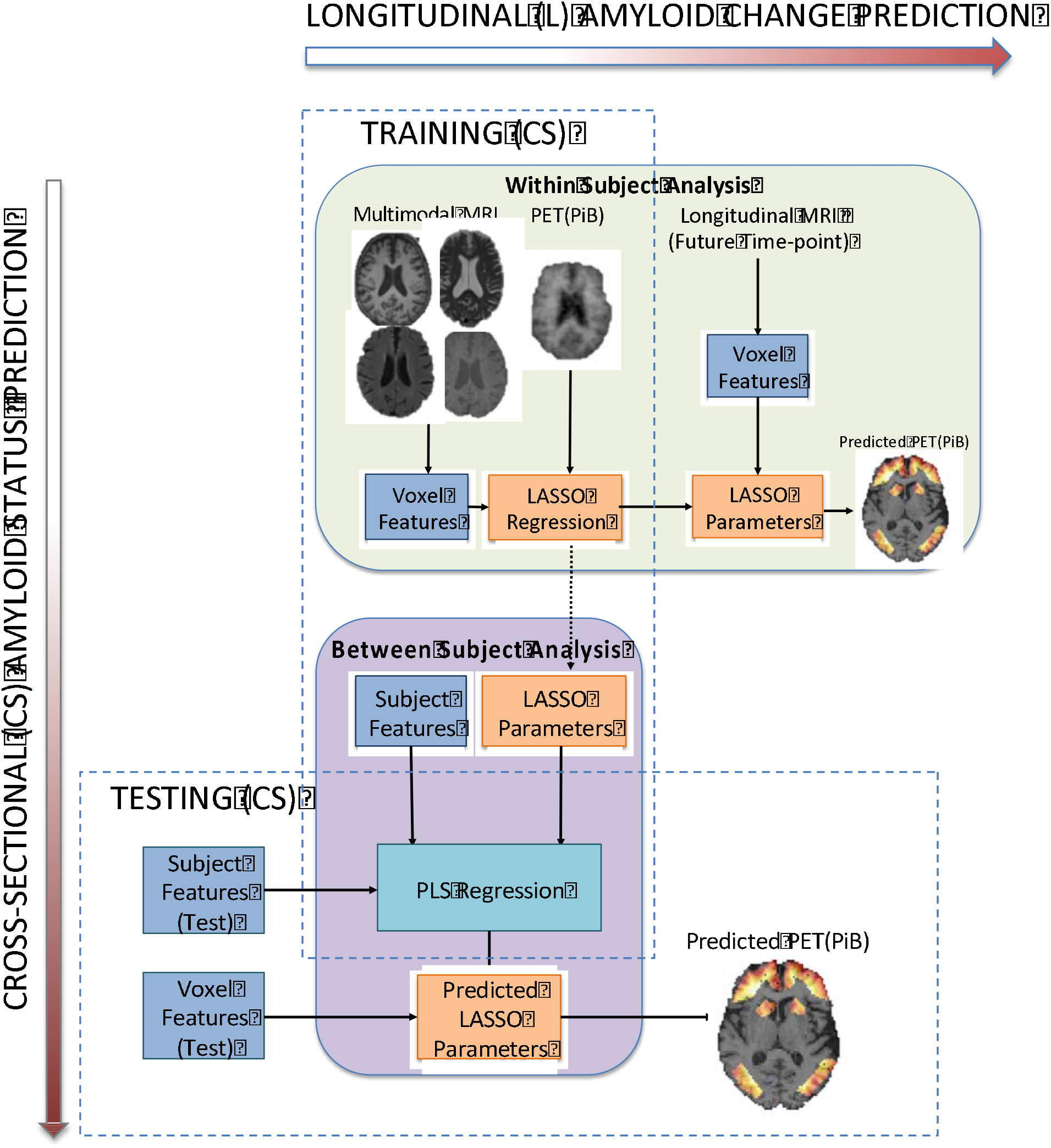
Overview of amyloid- β change prediction and amyloid- β status prediction

**Figure 2.**
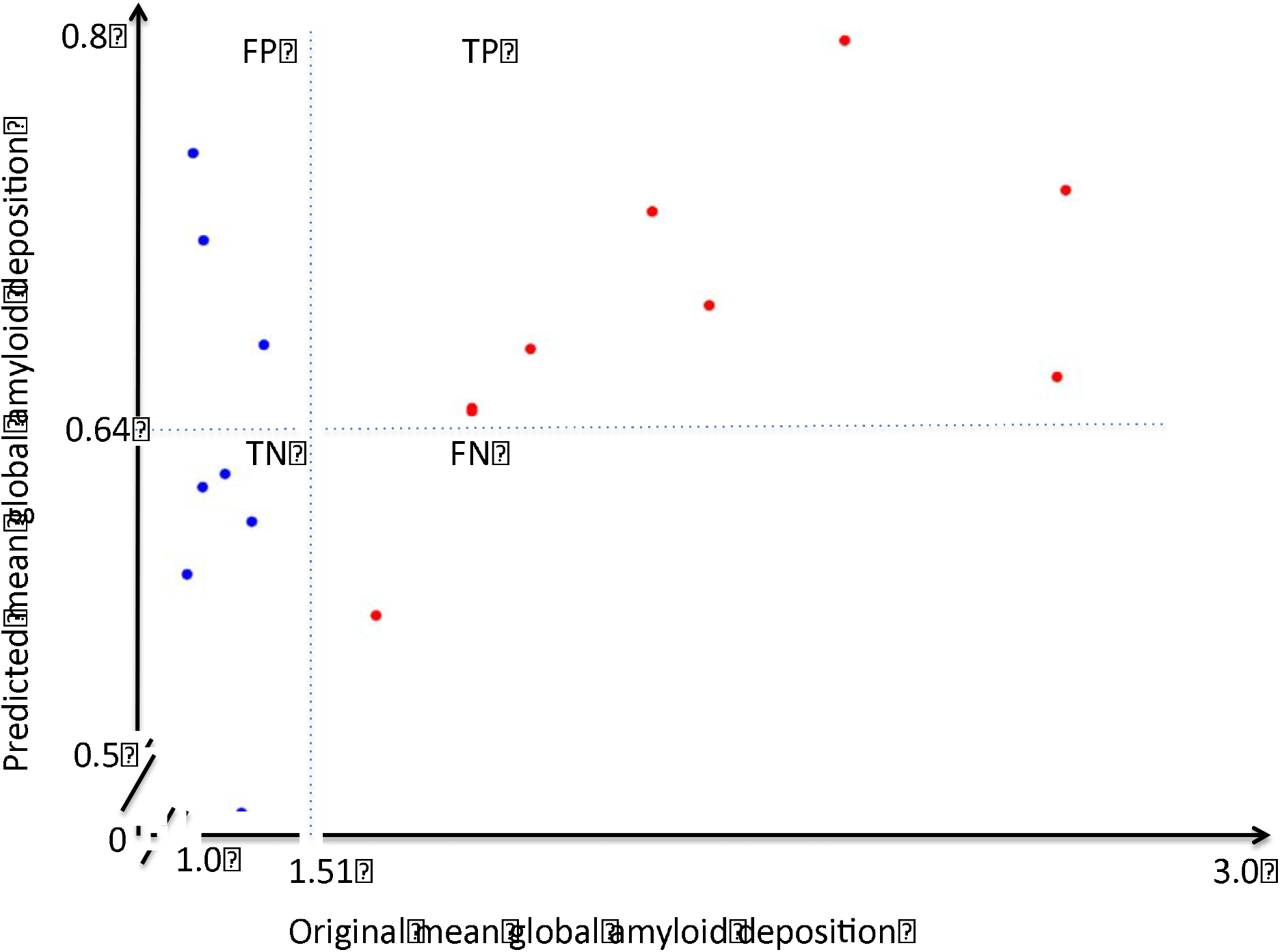
Mean global amyloid- β (within 6 ROIs) deposition in original and predicted for across subject amyloid- β status prediction using T1-weighted, T2-weighted FLAIR and SWI. Colors represent the ground truth: original PiB- (blue) original PiB+(red) (shown for one subset)

### Amyloid- β change prediction within subject

For predicting the increase in amyloid- β across two timepoints, T2-weighted imaging alone had the the best performance [For N=13 subjects: F-score: 83.33, accuracy = 84.62; AUC = 0.85 Spearman correlation coefficient = 0.81] (Table 2.).

## Discussion

The different MR imaging modalities carry complementary information about regional structural integrity. Several imaging features, from each of these modalities, were combined in a two-level approach (voxel-level and subject-level) to estimate amyloid- β status across subjects and estimate within-subject change in amyloid- β. Results of our analyses demonstrate that estimation of amyloid- β deposition could be successfully achieved using individual or a combination of MR modalities. In this approach, subject-level learning involved generalizing the LASSO parameters across subjects, hence inter-subject registration using template normalization – which can introduce variability – was not required. The approach predicts voxel-level amyloid- β, which can aid in both qualitative and quantitative analysis of regional amyloid- β burden.

For across subject amyloid- β status prediction, the combination of T1-weighted, T2- weighted FLAIR and SWI modalities were most predictive. This could be because the summary subject-features were most correlated with these modalities. The subject-level MR features like WMH and hippocampal volume are obtained from T1-weighted and T2-weighted FLAIR. The FP could be indicative of increasing regional amyloid- β burden although this finding still needs full validation. The PiB status prediction was better using both voxel-wise MR and summary subject-level features than using only summary subject-level measures. This suggests that there is information in MR signal at the voxel-level that is associated with amyloid- β accumulation, which drives the prediction of amyloid- β. The T2-weighted imaging alone had the highest prediction for amyloid- β change. T2 based contrast might capture the tissue changes in the gray matter (Diaz-de-Grenu et al., 2011). These changes might be helping in the detection of amyloid- β change over time.

Our key contribution in this work is to test whether there is a voxel-level association between MRI features and global amyloid- β deposition and show how this can be leveraged for amyloid- β prediction. Previous studies have explored the use of machine learning for cerebral amyloid- β prediction. Catell et al. (2016) used 3D gradient changes in amyloid- β imaging to classify amyloid- β status, their approach improving the detection of amyloid- β status from the PET images. A limited number of prior studies have used MR imaging to predict amyloid- β positivity. Ten et al. (2018) used a combination of features as predictors for amyloid- β status prediction (Ten Kate et al., 2018). They used subject demographics, cognitive variables, regional estimates of volume and cortical thickness from MRI, and *APOE* ε4 information along with machine learning classifier called support vector machine (SVM) with nested 10-fold crossvalidation to identify the best discriminating features between amyloid- β positive and amyloid- β negative groups. In our study however, we did not use any cognitive measures since we focused on preclinical biomarkers in cognitively normal older adults. To our knowledge, this is the first study that explores the use of voxel-level MR features to predict global amyloid- β deposition in cognitively normal subjects.

One limitation is our relatively small sample size. Although our current results suggest the voxel-level MR features can predict global amyloid- β status, we suspect with more modalities, more features, including asymmetry filters, and a larger set of images the learning would be much better. Deep learning approaches, such as convolution neural networks on large data sets is a promising future direction. This result demonstrates how voxel-level imaging data can be leveraged for prediction across individuals, and across time.

## Conclusions

In conclusion, this paper describes an approach for using image processing and pattern recognition techniques to predict amyloid- β deposition.

## Acknowledgments

Amyloid- β Pathology and Cognition in Normal Elderly (RF1 AG025516). Conflicts of interest: GE Healthcare holds a license agreement with the University of Pittsburgh based on the technology described in this manuscript. Drs. Klunk and Mathis are co-inventors of PiB and, as such, have a financial interest in this license agreement. GE Healthcare provided no grant support for this study and had no role in the design or interpretation of results or preparation of the manuscript. All other authors have no conflicts of interest.

