## Supplemental document for "Multimodal magnetic resonance imaging predicts regional amyloid- β burden in the brain"

*Author(s): Anusha Rangarajan^1^, Minjie Wu^2^, Naomi Joseph^3^, Helmet T. Karim^2^, Charles Laymon^1,4^, Dana Tudorascu^2,5,6^*, *Beth Snitz^7^, Annie Cohen^2^, Chester Mathis^3^, William Klunk^2,6^ and Howa­rd J. Aizenstein^1,2^*

*Affiliation(s):*

Departments of ^1^Bioengineering, ^2^Psychiatry, ^3^Chemical Engineering, ^4^Radiology, ^5^Biostatistics, ^6^Medicine, ^7^Neurology, University of Pittsburgh, Pittsburgh, PA, USA

*Corresponding author:

Howard Aizenstein, MD, PhD;

Charles F. Reynolds III and Ellen G. Detlefsen Endowed Chair of Geriatric Psychiatry

University of Pittsburgh, Department of Psychiatry,

Western Psychiatric Institute and Clinic, 3811 O’Hara Street,

**Supplemental Information**

**Feature Extraction**

**1. Voxel-level features**

The voxel-level features obtained from the MR images highlight distinguishable characteristics like intensity variations and changes. Features were extracted at the native resolution of each modality. Hence in addition to intensities and applying Gabor filters, local binary patterns and 3D gradients were obtained from T1-weighted imaging due to their high-resolution.

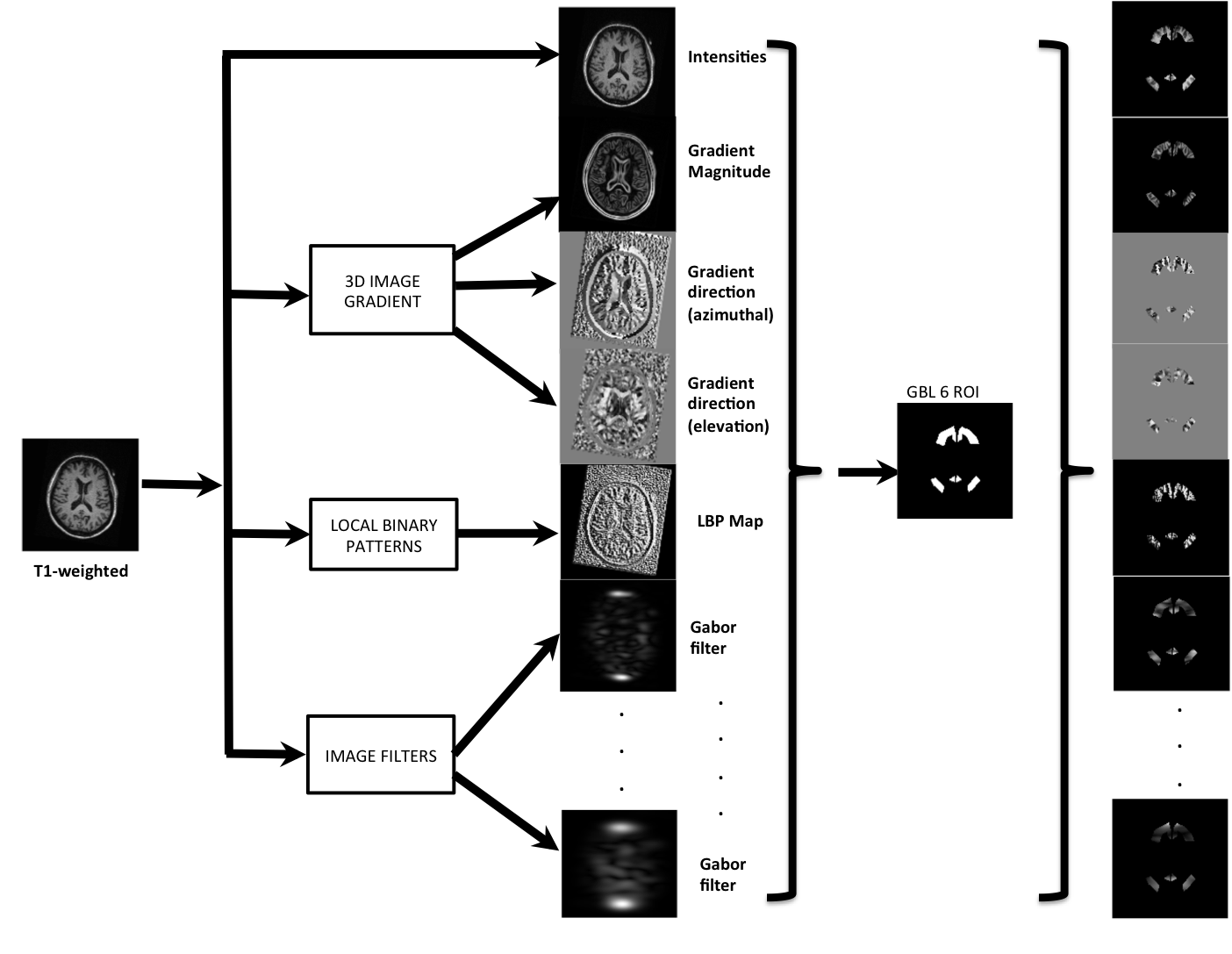

Figure 1 Voxel-level feature extraction for T1-weighted imaging: intensities, 3D gradient magnitude and directions, LBP map and Gabor filters.

*Gabor filters*

Gabor filters are linear frequency based filters, which highlight frequency contents in specific directions. Features constructed from responses of Gabor filters, Gabor features, have been successful in many computer vision and image processing applications face recognition, fingerprint matching. Gabor filter is the implementation of the Gabor transforms which is a short term Fourier transformation with Gaussian window for analysis in the spatial domain.

For obtaining the gabor residuals $u(x,y)$, convolution of an image $I(x,y)$ is done with 2D Gabor function $g(x,y)$ as represented by

Equation 1 Gabor residual u(x,y)

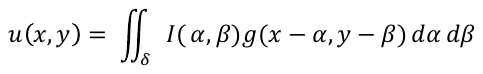

Where $x,y$ are set of image points and $\alpha,\beta$ are the integrals

Where $g(x,y)$ is the Gabor function and is given by

Equation 2 Gabor function g(x,y)

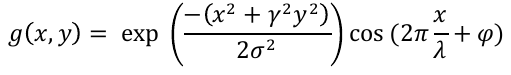

where $x=acos\theta+bsin\theta, y=-acos\theta+bsin\theta$

They are defined using different parameters (sizes 4 and orientation [45, 90, 180, 270 degrees]) each of these highlighting the change in intensities at different orientations.

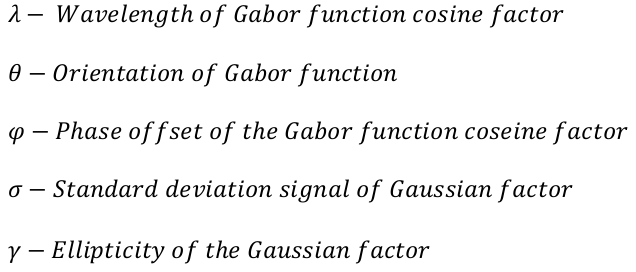

*3D Image Gradient:* Gradient filter applies the 3D Sobel spatial filter that highlights the edges, which are sudden changes in intensity. Sobel filter is a spatial high pass filter that allows visualizing rapid changes in intensities. Sobel filter is convolulved with the image to obtain the The Gx Gy and Gz are sobel filters in x-direction, y-direction and z-direction respectively.

$Gx= \left( \begin{matrix} -1 & 0 & 1 \\ -2 & 0 & 2 \\ -1 & 0 & 1 \end{matrix} \right)$ $, Gy= \left( \begin{matrix} -1 & -2 & -1 \\ 0 & 0 & 0 \\ 1 & 2 & 1 \end{matrix} \right)$ , $Gz= \left( \begin{matrix} -1 & 0 & 1 \\ -2 & 0 & 2 \\ -1 & 0 & 1 \end{matrix} \right)$

From these gradient vectors the gradient magnitude and angles (azimuthal and elevation) are computed.

$Gradient Magnitude= \sqrt{G_{x}^{2}+G_{y}^{2}+G_{z}^{2}}$

$Gradient angle_{azimuthal} = \tan^{-1} \left( \frac{G_{y}}{G_{x}} \right)*(\frac{180}{\pi})$

$Gradient angle_{elevation} = \tan^{-1} \left( \frac{G_{y}}{hypot(G_{x},G_{y})} \right)*(\frac{180}{\pi})$

Gradient elevation contains angles in degrees within the range [-90 90] measured between the radial line and the *x*-*y* plane while gradient azimuthal contains angles in degrees within the range [-180 180] measured between positive *x*-axis and the projection of the point on the *x*-*y* plane.

*1.4. Local Binary Patterns:* The local binary pattern is a texture-based feature which looks at its neighbors. Neighbors that are greater than the voxel value are coded as one and zero otherwise, and a binary value is generated (by going clockwise) – which is then converted to a decimal. Local binary patterns have been previously used for identifying these patterns in MRI (Maani, Kalra, & Yang, 2014; Oppedal, Eftestol, Engan, Beyer, & Aarsland, 2015). Fig. shows a 3x3 neighborhood, all values above 4 are considered 1, and rest are labeled 0. This generates a binary pattern clockwise as 11110000. The center value is then replaced with the decimal equivalent of the binary number. Based on the neighboring binary information spot, line edges/corners can be detected.

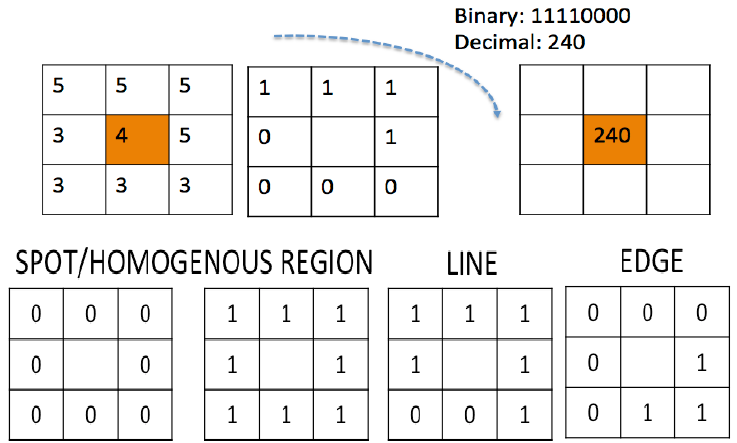

Figure 2 Local binary patterns

**2. Subject-level features**

*2.1. Participant demographics:* Demographic information included features such as age, sex, race, education, weight, and height (Tang et al., 2015)

*2.2. White Matter Hyperintensities (WMH):* White matter segmentation includes a fuzzy seed based segmentation(Wu et al., 2006) that identifies hyper intensities in the FLAIR that corresponds to the white matter lesions (Debette & Markus, 2010(Gorelick, 2011 #1813)). These are often seen in healthy older adults, and are more extensive in individuals with dementia. The automated WMH segmentation method is an iterative algorithm that involves an automated selection of “seeds” of possible WMH lesions and fuzzy connectedness, which clusters voxels based on their adjacency and affinity, to segment WMH lesions around the seeds. The fully automated WMH segmentation system was implemented in C++ and ITK. The total WMH volume divided by intracranial volume (ICV) was calculated as a marker of WMH burden.

*2.3. Hippocampal Volume:* Volume of the hippocampus is obtained using Automated Labelling Pathway (Dehmelt & Halpain) algorithm. ALP produces voxel counts for a large number of anatomically-defined brain regions, including all the Brodmann areas and subcortical structures, which have been hand-drawn on the atlas brain from the Montreal Neurological Institute (MNI).

2.4. Normalized gray matter and white matter: Gray matter and white matter probabilities were obtained using SPM12 (as described in 2.3. MRI preprocessing) and a binary image is obtained by using threshold of 0.6. Gray matter and white matter voxel counts were normalized by ICV.

**3. Stratified LOOCV for amyloid prediction across subjects**

The subject-level analysis involves a nested leave-one-out-cross validation (LOOCV) with stratified rule for maintain balanced PiB+ and PiB- subjects. Each subset consists of 18 subjects (9 PiB- and 9 PiB+). There are two LOOCV loops, outer LOOCV, for amyloid status prediction and inner LOOCV for obtaining the threshold used for prediction (figure 3). The outer LOOCV 17 subjects are used for training and 1 subject is left out. From 17 subjects, once again LOOCV is performed for predicting a threshold using linear regression. In the inner LOOCV, 16 subjects are used for training the PLS model and 1 subject is left out. Using subject features of left out test data and trained PLS model LASSO parameters are predicted. Predicted LASSO parameters are fit on the voxel level features from the test subject to obtain voxel-level amyloid prediction. This is repeated for all 17 subjects as test data and regression model is fit between mean original and predicted amyloid. Linear regression model acts as mapping from the original amyloid data to the predicted data, by means of which the threshold is obtained. The voxel-level amyloid is predicted on the outer test data from PLS model (obtained on all 17 subjects) and the respective subject-level and voxel-level features.

The amyloid voxel-prediction through the subject-level learning method described above is shown in figure 4. (Note: The original amyloid PET (PiB) images are also normalized from 0 to 1 for representation). The AUC plot for same subset (shown in the figure 2. in main manuscript) is given in figure 5.

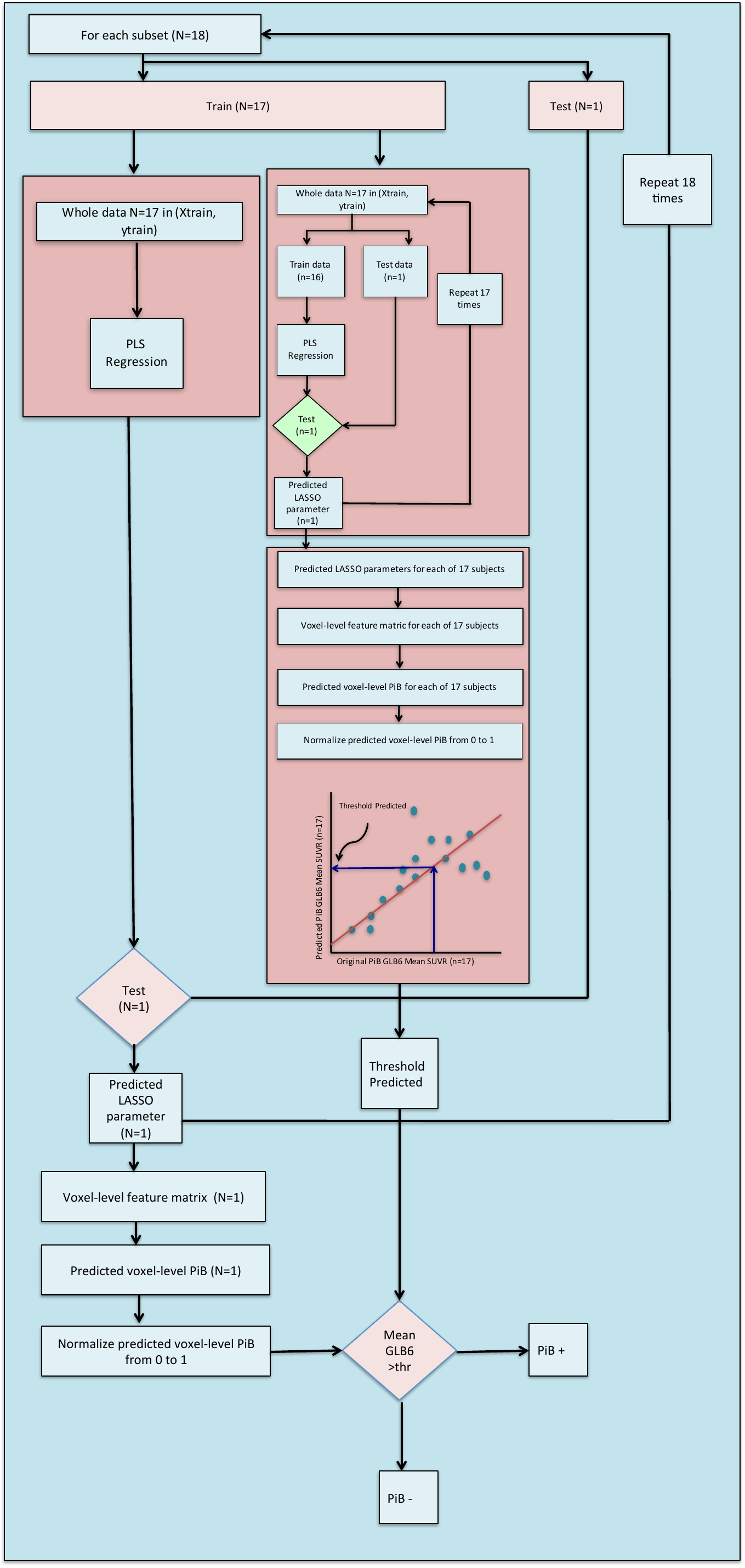

Figure 3 Subject-level analysis with nested LOOCV

Figure 4 Amyloid prediction voxel-level for PiB + and PiB- subejcts using T1-weighted, T2-FLAIR and SWI modality combination

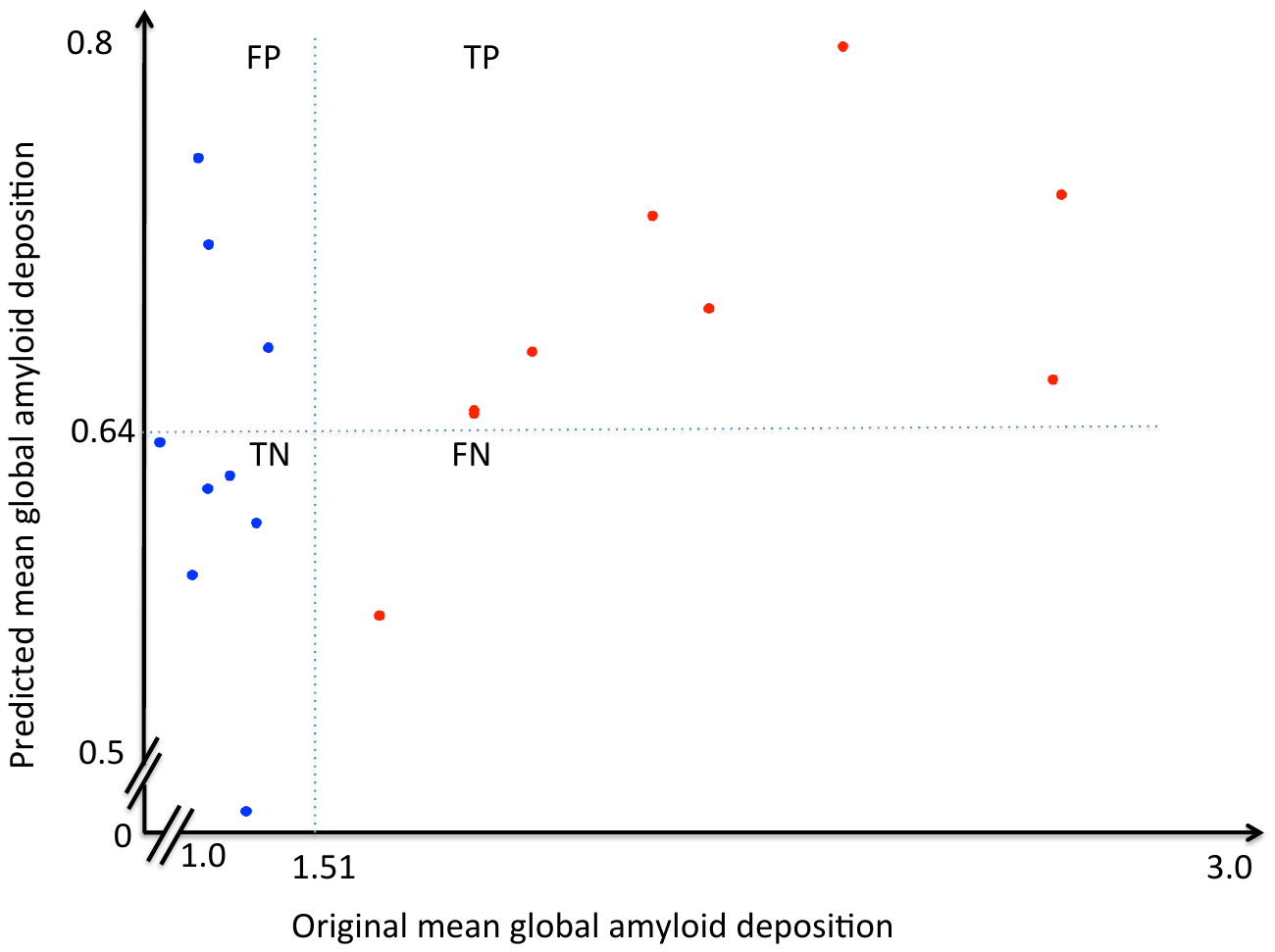

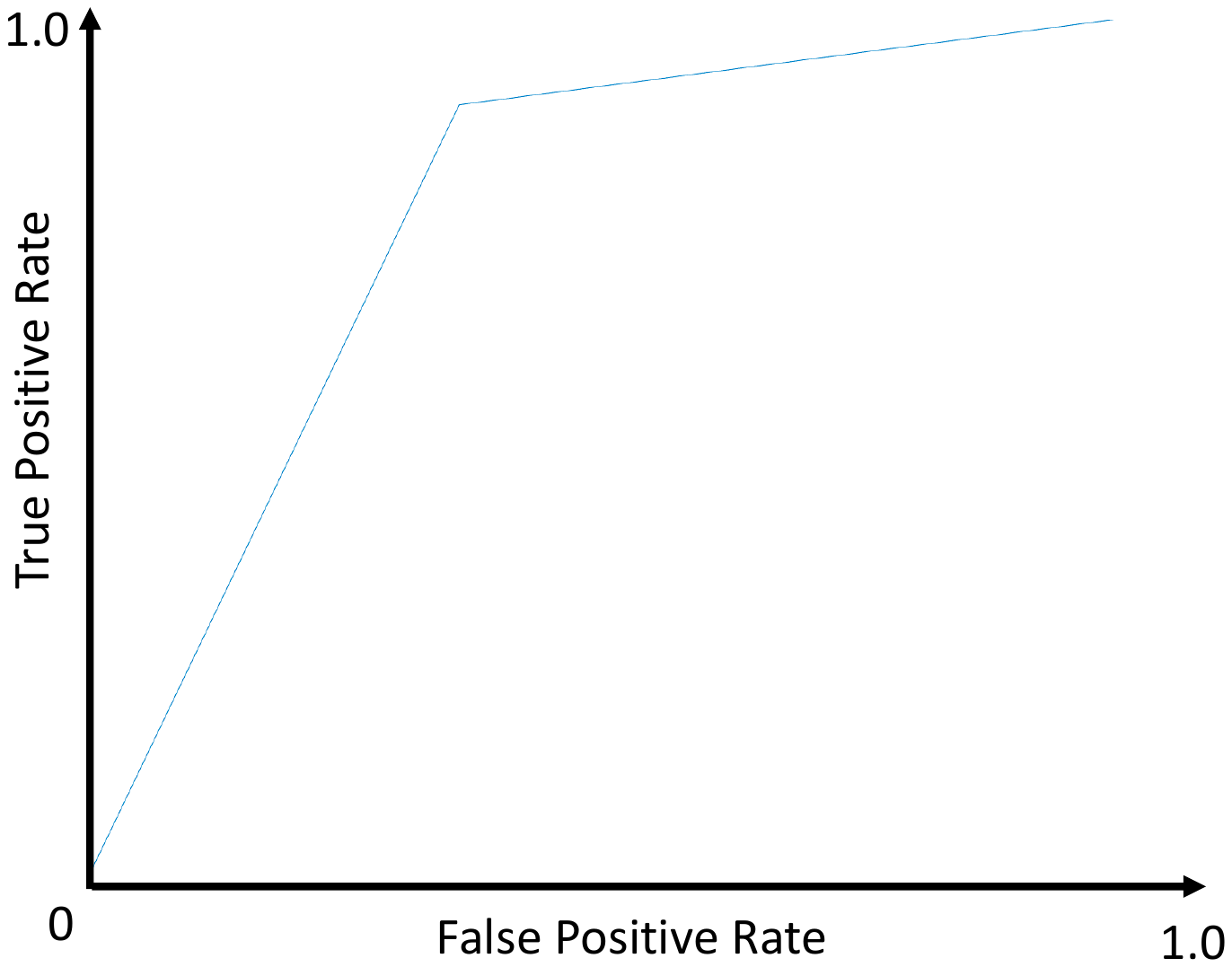

Figure 5. (Left) Mean global amyloid (within 6 ROIs) deposition in original and predicted for across subject amyloid status prediction. Colors represent the ground truth: original PiB- (blue) original PiB+(red) (shown for one subset);(Right) AUC plot for the subset
